# Modeling Paralinguistic Features for Speech Neuroprostheses: A Systematic Review

**DOI:** 10.64898/2026.10.11.758333

**Authors:** Manu Siddharth Yenikapati, Soham Deshpande, Shiv P Desai, Donovan D Santine, Aryan K Roghani

## Abstract

Recent advances in brain-computer interface (BCI) technology have brought speech neuroprostheses closer to restoring naturalistic communication for individuals with severe motor or speech impairments. Although current systems can decode phonemic and lexical information from cortical activity, they often fail to capture paralinguistic dimensions such as intonation, prosody, loudness, and affect that are essential for conveying emotion and intent. This review synthesizes emerging research on modeling and incorporating paralinguistic features into speech neuroprosthetic design. Foundational studies have mapped the cortical organization of speech production, identifying articulatory somatotopy within the sensorimotor cortex and functional subdivisions across planning, phonological, and semantic representations. More recent work has characterized expressive encoding within the ventral precentral and supplementary motor areas, while peripheral and multimodal sensing approaches have demonstrated promising accuracy in decoding emotional states. Collectively, these findings signal a shift toward closed-loop neuroprosthetic systems that integrate affective modulation with linguistic decoding, enabling restoration of speech that is not only intelligible but expressively human.

## Introduction

Rapidly advancing development in the fields of artificial intelligence and language learning models (LLMs) has integrated with the development of brain-computer interface (BCI) technology^1,2^, allowing neural signals to serve as input for algorithms that can predict expressive output^3,4^. BCI research has already led to a multitude of innovations, including prosthetics and disorder monitoring systems^5,6^. Early breakthroughs have already suggested that lexical information, including phonemes, words, and sentences, can be decoded enough to support functional communication, marking a major step towards speech restoration for those suffering from lost motor control^7^.

Due to various causes spanning from Amyotrophic Lateral Sclerosis (ALS)^8^ to brainstem stroke^9^, many people exhibit dysarthria, the difficulty of producing clear and articulate speech, or in some cases, anarthria, the near-complete inability to produce clear and articulate speech^10^. In either case, the ability to produce naturalistic speech is drastically lowered, which can dramatically decrease one’s quality of life^11^. In recent years, there has been a focus on developing BCI technologies that utilize LLMs to predict speech based on brain input, known as speech neuroprostheses^4,12^. However, the current focus has been on developing solely textual interfaces based on lexical decoding^10,13,14^. Despite the obvious benefits of this technology, restoring naturalistic communication in patients who exhibit dysarthria or anarthria requires the ability to locate the physical markers of paralinguistic features^15^ such as intonation, loudness, prosody^16,17^, and affect that have an effect on expressivity apart from lexical expressions^18,19^.

The rapid progress of BCI technologies for naturalistic speech restoration^20,21^ highlights the growing need for a comprehensive synthesis of current developments in paralinguistic modeling to inform new directions for research. While numerous studies have advanced lexical and semantic decoding from cortical activity^22,23^, efforts to capture paralinguistic dimension remain sparse^15^, and no recent systematic review has captured these developments. To address this gap, this review consolidates emerging research on the neural encoding and modeling of expressive speech features and their integration into speech neuroprostheses, with the goal of guiding future systems to restore communication that is not only accurate but authentic.

## Method

### Search Strategy

This systematic review was conducted in accordance with the latest guidelines of the Preferred Reporting Items for Systematic Reviews and Meta-Analyses (PRISMA)^24^. The search was conducted on October 24, 2025 on Google Scholar and PubMed using the following two queries:

1. ((“neuroprosthesis” AND (“paralinguistic” OR “naturalistic communication”) AND “speech”)
2. (“prosody” OR “intonation” OR “affective speech” OR “emotional prosody”) AND (“BCI” OR “brain–computer interface” OR “neural decoding”)

On Scopus, the search was conducted using the following query:

1. “TITLE-ABS-KEY (speech AND (neuroprosthesis OR “brain-computer interface”)) AND TITLE-ABS-KEY (paralinguistic OR prosody OR intonation OR affect*)”

A secondary verification search was conducted on November 5, 2025 to ensure no newly indexed studies were missed, yielding the same results as the initial search.

Given the narrow and emerging nature of paralinguistic modeling in speech BCIs, broader queries resulted in a high proportion of irrelevant lexical-only decoding studies. Pilot searches indicated that expanding beyond the two final query structures increased noise without improving recall. Therefore, two high-specificity queries were selected to balance sensitivity and relevance.

### Selection Criteria

One author (M.Y) performed the search and reviewed all articles using the aforementioned databases.

The inclusion criteria were as follows. (1) Studies that had an express focus on paralinguistic features. (2) Studies that had a focus on the applicability of their research to BCI or speech neuroprosthesis development.

The exclusion criteria were as follows. (1) Review articles. (2) Studies that focused only on semantic or lexical decoding. (3) Articles that were not available in English.

### Categorization and Refinement

While reviewing all articles gathered from the aforementioned query, the articles were all sorted into seven distinct categories, as follows:

1. Literature Reviews of Speech and Communication.
2. Empirical Studies on Neural Encoding and Decoding of Speech.
3. Applied Speech Neuroprosthesis.
4. Computational Advances in Neural Speech Modeling.
5. Multimodal Approaches to Speech and Voice Modeling.
6. Ethical Dimensions of Neural Speech Interfaces.
7. Systematic Reviews of Neural Speech Interface Research.

Categories were intentionally broad to capture the full ecosystem of neural speech interface research. Although some categories were excluded at later stages, their inclusion during initial screening ensured that paralinguistic-focused studies were not prematurely discarded due to disciplinary overlap.

Studies were initially screened by the title and abstract, then by full-text review. Categories (1) and (7) were excluded due to exclusion criterion (1) (Review articles). Category (6) was excluded because it did not have a focus on paralinguistic features. Articles assigned to categories (2), (3), (4), and (5) were included on a case-by-case basis according to the inclusion criteria defined above. All abstracts of articles in these categories were reviewed by another author (S.D) to finalize the group of included articles (κ = 0.6846, indicating substantial agreement^25^). Disagreements were resolved via mutual deliberation in line with the previously outlined criteria. The decision-making process is illustrated below (Figure 1).

**Figure 1.**
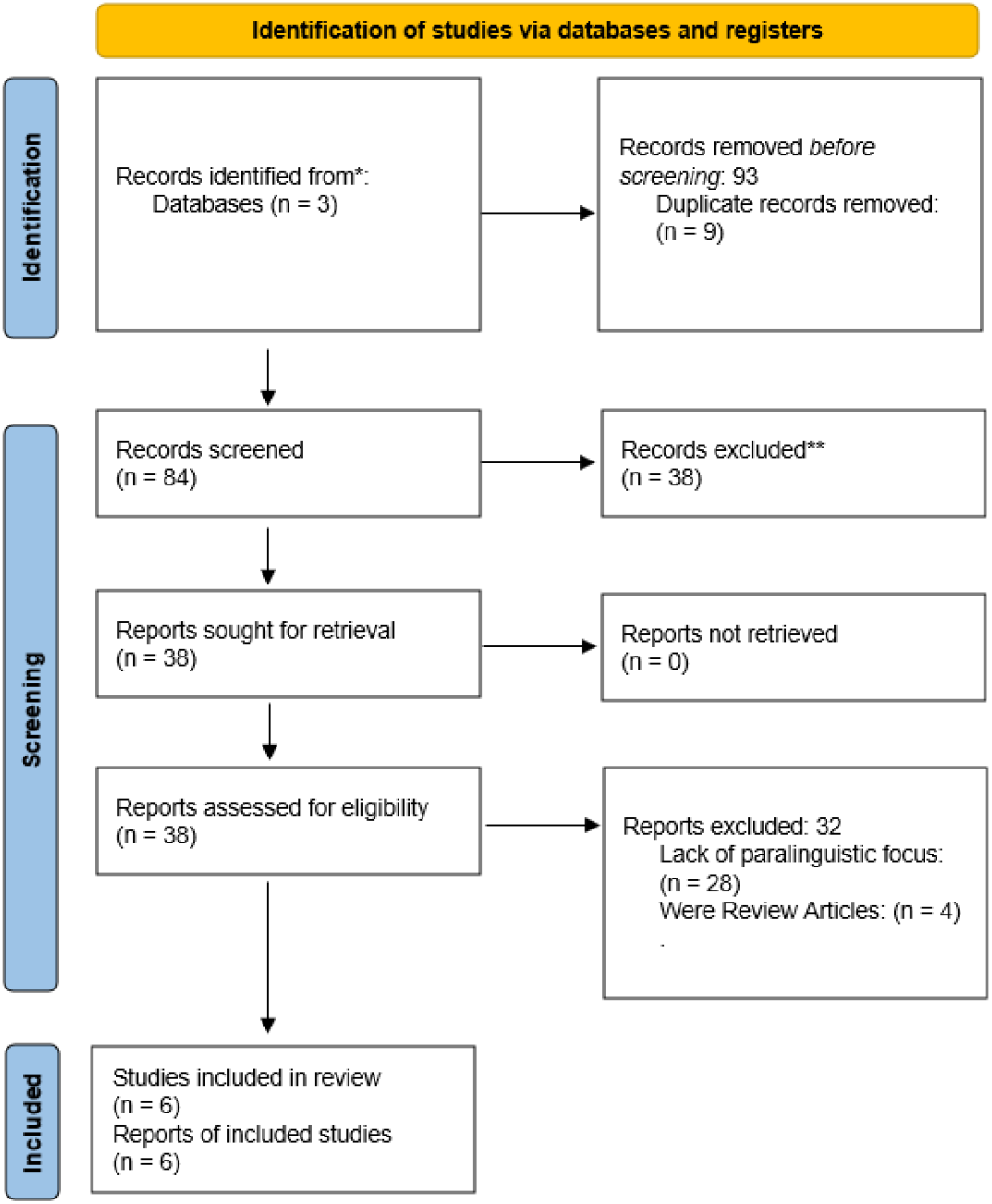
PRISMA Flowchart Illustrating Article Selection Process. Note: PRISMA flow diagram illustrating the systematic search and selection process for studies examining paralinguistic feature modeling in speech neuroprostheses. Records were identified through database searches of PubMed, Scopus, and Google Scholar, followed by duplicate removal, title and abstract screening, full-text review, and final inclusion based on predefined eligibility criteria. Reasons for exclusion at each stage are indicated.

**Figure 2.**
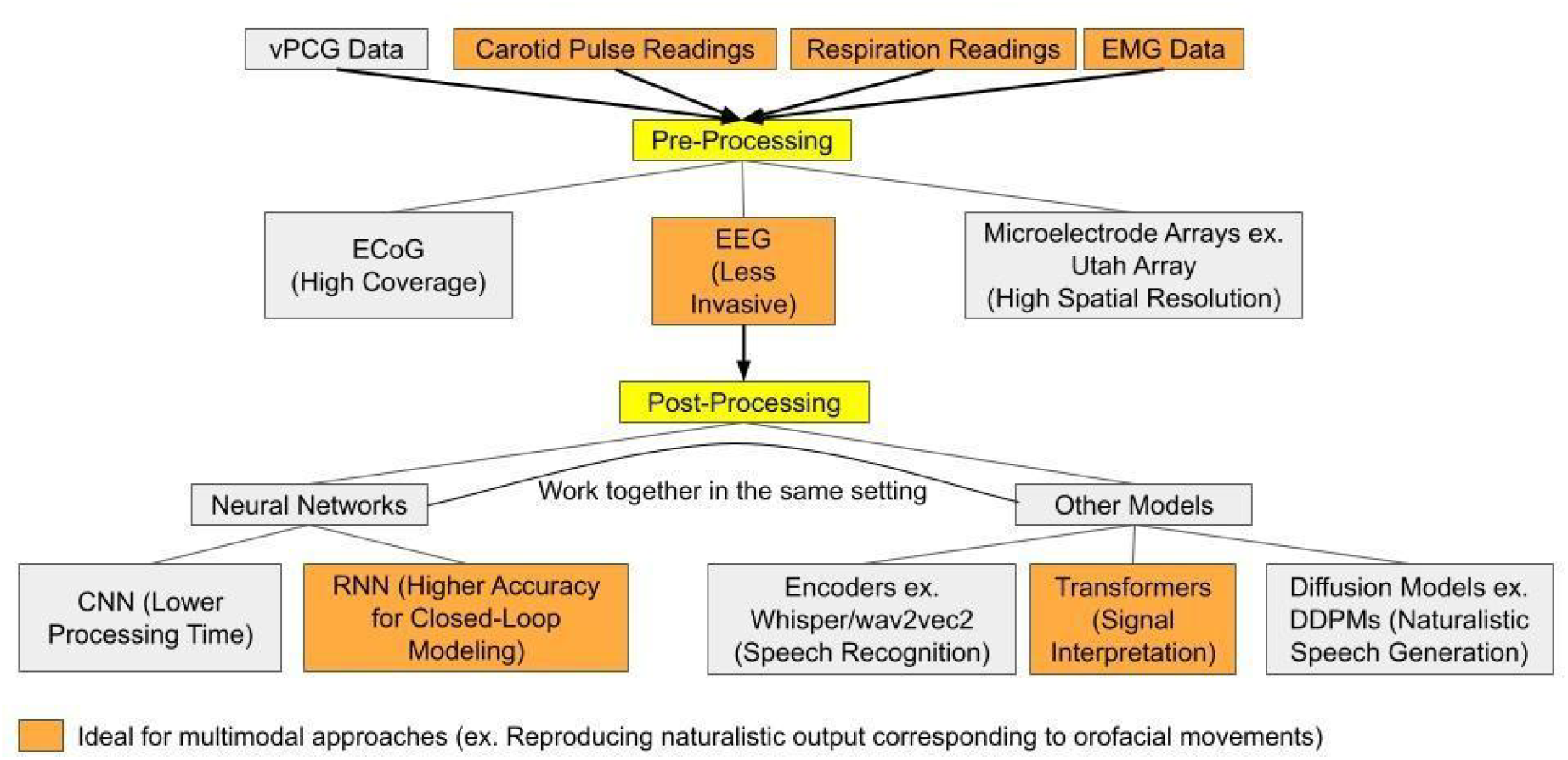
Signal Processing Pipeline for Paralinguistic Feature Modeling in Speech Neuroprostheses. Note: Conceptual pipeline illustrating how neural and peripheral signals can be transformed into continuous paralinguistic trajectories within a closed-loop speech neuroprosthesis

Due to the novel nature of the research question at hand, most studies regarding applications of speech neuroprostheses in paralinguistic modes are relatively new and have not had the opportunity to be published. As a result, four of the six articles were preprints at the time of writing. It has been contested that the lack of peer review renders these sources unreliable, especially when conducting systematic reviews^26^. However, the inclusion of preprints within systematic reviews has been justified via the observed lack of significant reduction in quality that comes with preprints^27–29^.

## Results

An initial search on October 24, 2025, yielded 93 total articles, of which 9 were duplicates, and 46 were excluded either because they were review articles or because they did not focus on speech neuroprosthesis/BCI technology. Of the remaining 38 articles, 32 were excluded due to a lack of primary focus on paralinguistic features instead of lexical encoding or decoding. The final review included six studies (2023-2025) examining the ways in which paralinguistic features could be modeled via BCIs to develop novel speech neuroprosthesis technology (Table 1). Studies used one to five participants and utilized various modalities to record neural activity, such as electrocorticography (ECoG), electroencephalography (EEG), microelectrode arrays such as the Utah Array, and carotid pulse readings. Various neural network models were used to decode neural data, such as recurrent neural networks (RNNs), convolutional neural networks (CNNs), and multinomial logistic regressions.

**Table 1.**
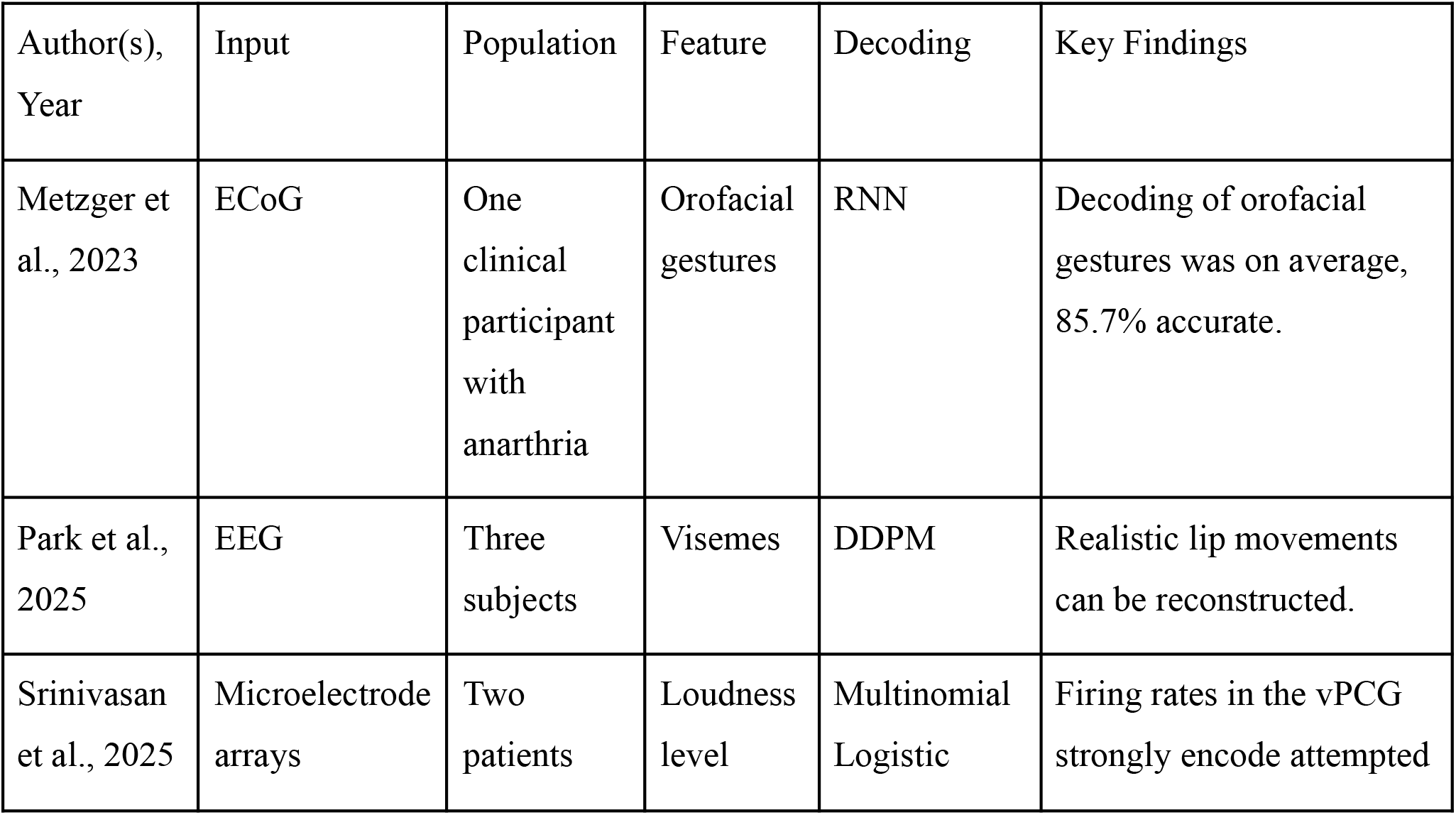
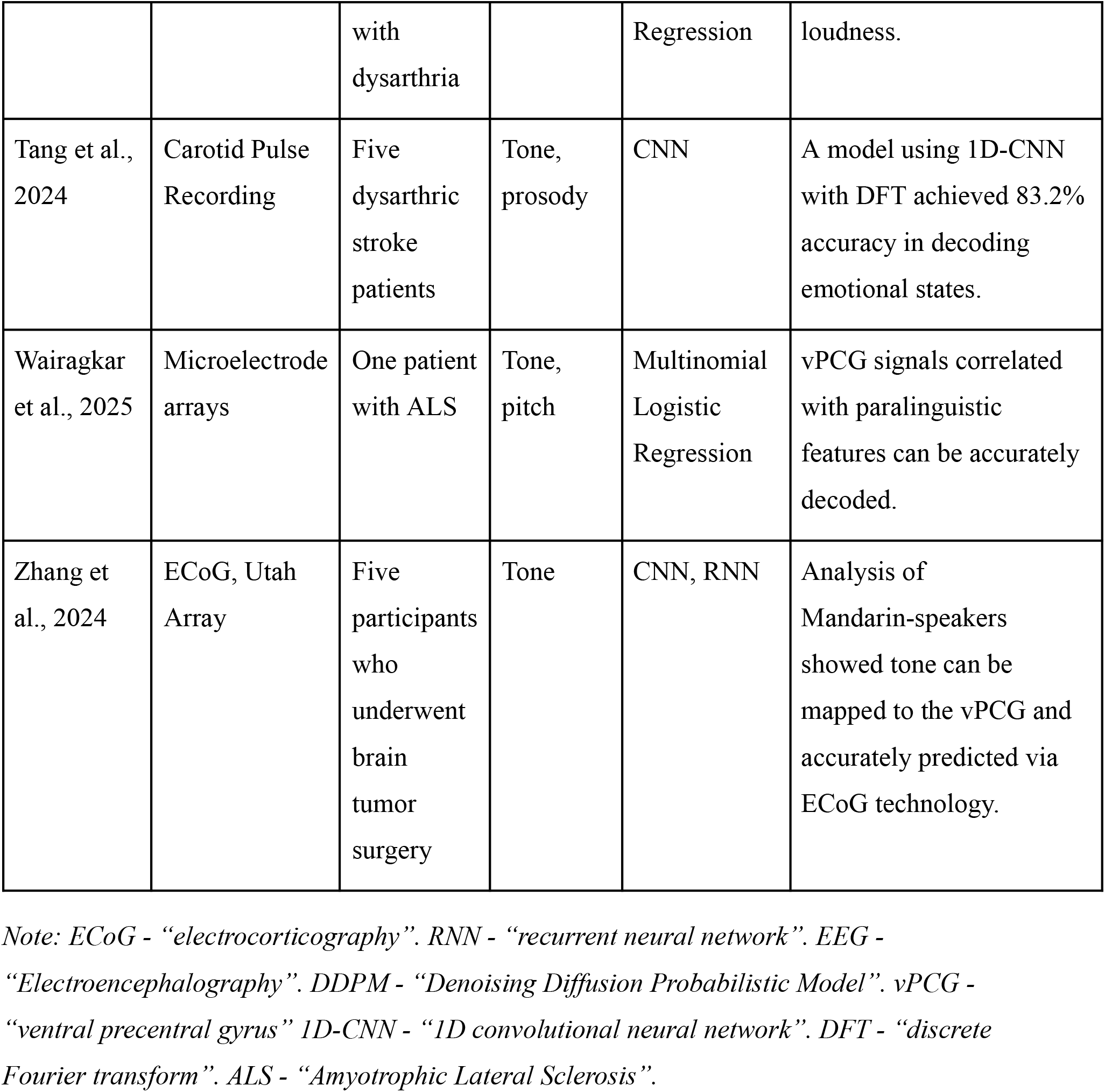
Existing Studies in Paralinguistic Modeling for Speech Neuroprostheses.

## Discussion

The key findings of the six selected studies could broadly be categorized into three groups based on the stage of speech neuroprosthesis development a certain finding offers insight into.

### 1. Data Collection of Paralinguistic Features

Paralinguistic features vary greatly in form; expressions such as orofacial gestures, intonation, and loudness may require different methods of data collection^15^. Data pre-processing must be aligned towards the targeted feature. ECoG, EEG, and Utah Arrays are most common for measuring vPCG signals that appear to encode information such as intonation^30,31^ and loudness ^32^, however other biosignals, such as carotid pulse readings^33^, are increasingly used as a measure of affect in attempted speech^15^.

Additionally, novel techniques are being used to develop predictive machine learning algorithms that can be used in conjunction with a BCI to produce output that resembles naturalistic speech^10,34^. Historically, computational paralinguistic systems relied on conventional classifiers, but modern architectures increasingly employ RNNs and CNNs^35,36^. Especially in closed-loop environments, RNNs seem to be more suited to the task at hand given their ability to process real-time sequential data^37,38^. Additionally, emerging architectures such as transformers^39–41^, diffusion models^42^, and other speech encoders including Whisper and wav2vec2^43,44^ have demonstrated the capability to capture subtle acoustic and temporal patterns to decode expressive features from neural data. Hence, these types of models can generate trajectories of pitch and amplitude, as well as learn from large unlabeled datasets, making them well-suited to reconstruct naturalistic prosody, especially when combined with neural network models such as RNNs and CNNs^45^.

### 2. Neural Encoding of Paralinguistic Features

In order to develop a speech neuroprosthesis that is capable of modeling paralinguistic features, it is first necessary to understand the physical impetuses that cause expressive communication. Recent research suggests that the ventral precentral gyrus (vPCG) appears to play a central role in paralinguistic expression^31,32^. Since the vPCG plays a crucial role in integrating fine-grained motor planning^46,47^ for articulators like the lips, jaw, and larynx with modulation of prosody^32^, this is associated with modulation of pitch and loudness in real time^48^. A machine learning algorithm that analyzed vPCG signals via demixed principal component analysis revealed that a speech BCI trained on this achieved a loudness classification accuracy of 94%^32^. A study of Mandarin speakers, who are known to express greater tonal variation than English counterparts^49^, found that tone could be cortically represented in the vPCG with area under curve scores (AUCs) 0.96-0.99^30^. In addition to loudness and tone, orofacial movements were found to demonstrate similarly strong associations in the sensorimotor cortex (SMC)^50^. The high accuracy of existing studies that have attempted to model paralinguistic expression biologically strengthens the feasibility of a speech neuroprosthesis that can incorporate these insights into its design.

### 3. Decoding of vPCG Data

Developing speech BCI technologies also necessitates the ability to decode vPCG data to make sense of neural input. Despite the dominant focus on studying open-loop systems^51^, a device that attempts to restore naturalistic communication requires the ability to incorporate data in a closed-loop system^52^. Such systems have been tested on vPCG data collected via ECoGs and Utah Arrays that display affectual decoding accuracies of approximately 80%^31^ and tonal decoding accuracies of approximately 46%^30^. EEG data also revealed that closed-loop systems for imagined viseme decoding had a viseme error rate of approximately 77.96%^53^.

Additionally, the fusion of multimodal outputs will most likely be essential for closed-loop expressive decoding^54,55^. The blend of neural activation with physiological signals like respiration and electromyograms (EMGs) could improve emotional clarity and stabilize predictions^56,57^. When paired with other advanced sequence models like transformers and DDPMs that can synthesize continuous paralinguistic trajectories, this multimodal fusion is even more impactful^58^.

### 4. Clinical Relevance and Translational Implications

The clinical relevance of incorporating paralinguistic features into speech neuroprostheses is most evident in patient populations for whom lexical decoding alone fails to restore functional communication, including individuals with anarthria or severe dysarthria due to amyotrophic lateral sclerosis, brainstem stroke, or other neurological injury^8,9,19^. In these conditions, affective intent and expressive modulation often remain intact at the neural level despite impaired motor execution. Paralinguistic features such as prosody, loudness, and emotional tone are therefore not merely aesthetic components of speech but convey clinically meaningful information^16,18^ related to intent, urgency, and social pragmatics, which are critical for safe and effective communication in daily and clinical interactions.

Restoring expressive modulation has the potential to improve quality of life and psychosocial outcomes beyond intelligibility alone. Conventional augmentative and alternative communication systems and text-based speech BCIs frequently impose emotionally neutral output, which can contribute to frustration, reduced agency, and social withdrawal. By enabling user-driven modulation of expressive features, paralinguistic-aware neuroprostheses may better support authentic self-expression, interpersonal engagement, and patient autonomy. These outcomes align with functional communication frameworks in neurorehabilitation, which emphasize participation and communicative effectiveness rather than accuracy in controlled laboratory tasks.

From a translational perspective, clinical feasibility depends on integrating paralinguistic decoding without substantially increasing surgical burden or system complexity. Converging evidence that the ventral precentral gyrus encodes multiple expressive dimensions^30–32^ suggests that paralinguistic features may be incorporated into existing electrode configurations used for articulatory decoding. Additionally, closed-loop feedback mechanisms are likely essential for clinical adoption^51,54^, allowing users to iteratively refine expressive intent through real-time auditory or visual feedback.

## Limitations

Despite its meticulous selection and review process, there exist several limitations of both this review and the supporting literature that warrant further consideration:

### 1. Sparse Data Sets

Only six studies were selected after the rigorous article screening process, and each of those studies studied between one and five participants, for a total of seventeen participants divided across all six studies. This emphasizes possible imperfections within the studies, underscoring the need for more data acquisition regarding paralinguistic features.

### 2. Imperfect Evaluation Metrics

Most of the selected studies quantified positive outcomes via low error rates, which do not meet the standard of mimicking naturalistic speech, as naturalistic paralinguistic output appears in great variety rather than in neatly categorized buckets that can be tested for accuracy.

### 3. Technology Constraints

Neural decoding of expressivity is limited by the signal-to-noise ratio (SNR)^59^ and cortical sampling constraints^60^, which is a limiting factor for analyzing the accuracy of decoding mechanisms and developing better technology.

### 4. Task Variability Across Studies

The reviewed studies offered many different decoding techniques for paralinguistic features, ranging from tone classification and loudness estimation to orofacial gestures and affect recognition. Although effective in their respective ways, the tasks vary in complexity and representation of neural data, making it difficult to compare results and truly generalize findings for neural decoding and expressive communication. As an example, decoding tone requires heavy fine-grained pitch modulation^61^, whereas affect recognition blends autonomic and motor cues^62,63^, and this variability limits the ability to draw broader conclusions regarding the performance of paralinguistic decoding.

## Future Directions

This systematic review highlights the necessity of further research regarding paralinguistic feature modeling in the production of speech BCIs. Future directions for possible research include:

### 1. Development of Multimodal Datasets

To address the low sample sizes of both this systematic review and of the selected underlying studies, the development of multimodal datasets that collect data regarding paralinguistic features such as intonation, loudness, orofacial gestures, and so forth, could be used to both train machine learning algorithms and reinforce the conclusions of the six studies analyzed.

### 2. Focus on Closed-Loop Systems

Despite the growing interest in modeling speech in a closed-loop setting, most studies are still conducted in an open-loop environment^51^. A focus on developing more accurate algorithms to predict expressive output in real-time could be the next step in developing more accurate and natural speech neuroprostheses.

### 3. Neural-Avatar Coupling for Multimodal Communication

The development of avatars based on multimodal neural input could help with data collection via allowing researchers in clinical settings to collect large volumes of data via ECoG or EEG readings^64^. Currently, avatars are used primarily to model one feature, as multimodal modeling might make response times too slow to operate in a closed-loop environment^15^, but tackling this issue could help advance algorithm development.

### 4. Accuracy Metric Enhancement

As outlined above, the usage of error rates as a primary measure of algorithmic success fails to reproduce speech at the naturalistic level. Using continuous-time approximations rather than discrete measures of paralinguistic features in clinical studies could yield more accurate estimates of the accuracy of machine learning algorithms^65^ and train their output to be continuous^66^, which is more representative of naturalistic communication.

### 5. Increased Targeting of Paralinguistic Features

The relatively low number of articles that met the selection criteria points to the fact that the modeling of paralinguistic features in speech neuroprostheses is under-researched, and this review points to the potential significance of bolstering research efforts in this direction to restore naturalistic communication within patients experiencing certain forms of communication barriers.

## Conclusion

The findings of this systematic review indicate the possibility of a pivotal transition in the field of speech neuroprosthetics, from decoding lexical and phonemic content to modeling the expressive, affective, and prosodic dimensions of human communication. Although current research remains limited in scale, with small participant pools and a narrow range of modalities, the reviewed studies collectively demonstrate that paralinguistic features such as tone, loudness, prosody, and orofacial gestures can be robustly encoded and decoded from cortical activity, particularly within the vPCG. These insights underscore the biological feasibility of neuroprosthetic systems capable of restoring speech that conveys not only words but emotional nuance and intent.

The integration of advanced neural decoding techniques, including recurrent and convolutional neural networks, has enabled preliminary success in reconstructing paralinguistic signals from both invasive and noninvasive modalities. However, achieving naturalistic communication will require a combination of multimodal data acquisition, improved signal fidelity, and the development of closed-loop architectures that adaptively respond to user intent in real time.

Furthermore, further research should move beyond categorical classification metrics towards continuous, context-sensitive measures of expressive quality to more faithfully approximate natural speech.

Ultimately, the path forward lies in uniting computational modeling, neuroscience, and affective science to design neuroprostheses that restore not just speech intelligibility, but the full expressive richness of human interaction. By embedding paralinguistic understanding into the next generation of speech BCIs, researchers move closer to bridging the gap between technological articulation and truly naturalistic communication.

